# Transgeneratonal inheritance of ethanol preference is caused by maternal NPF repression

**DOI:** 10.1101/536599

**Authors:** Julianna Bozler, Balint Z Kacsoh, Giovanni Bosco

## Abstract

Rapid or even anticipatory adaptation to environmental conditions can provide a decisive fitness advantage to an organism. The memory of recurring conditions could also benefit future generations, however neuronally-encoded behavior isn’t thought to be inherited across generations. We tested the possibility that environmentally triggered modifications could allow “memory” of parental experiences to be inherited. In *Drosophila melanogaster*, exposure to predatory wasps leads to inheritance of a predisposition for ethanol-rich food for five generations. Inhibition of Neuropeptide-F (NPF) activates germline caspases required for transgenerational ethanol preference. Further, inheritance of low NPF expression in specific regions of F_1_ brains is required for the transmission of this food preference: A maternally derived *NPF* locus is necessary for this phenomenon, implicating a maternal epigenetic mechanism of NPF-repression. Given the conserved signaling functions of NPF and its mammalian NPY homolog in drug and alcohol disorders, these observations raise the intriguing possibility of NPY-related transgenerational effects in humans.

## Introduction

To what extent is personality and behavior predetermined at birth? Philosophers and scientists alike have struggled with this question, and many have settled on the *tabula rasa*, or *blank slate* perspective. This long-standing notion posits we are without form or direction until our individual experiences shape us. Over the past decades however, evidence has accumulated that suggests parental environment can have significant phenotypic consequences on the next generation, thus eroding this notion of a blank slate. The Dutch Hunger Winter Study was one of the first documented examples of ancestral experiences influencing subsequent generations. Children conceived in the Netherlands during the World War II blockade, and ensuing famine, had higher rates of obesity and diabetes (Heijmans et al., 2008; Schulz, 2010; Stein, Susser, Saenger, & Marolla, 1975). More recent studies have found that neurological and mental health conditions also appear to have persistent impact on the next generations (Yeshurun & Hannan, 2018). Further, risk factors for children of Holocaust survivors, such as reduced cortisol sensitivity has been linked to methylation state of the glucocorticoid receptor promoter, and increased methylation in offspring was associated with paternal diagnosis of posttraumatic stress disorder(Yehuda et al., 2014).

Studied largely in the public health context, there are limited examples of environmental inheritance that can be experimentally tested. Genetic model systems thus are indispensable for understanding molecular mechanisms of causation. For example, male mice trained to associate fear with an odor-transmitted sensitivity of this odor to their sons. In this instance, researchers concluded that offspring possessed an increased abundance of sensory neurons specific to the same odor their fathers were trained to fear (Dias & Ressler, 2014). Similarly, environmental enrichment activities can ameliorate behavioral defects of mutant mice defective in long-term potentiation and memory, and this behavioral rescue is heritable to the next generation through the activation of an otherwise latent p38 signaling cascade (Arai, Li, Hartley, & Feig, 2009). Parental exposure to toxins and nutritional challenges also can change germline information, affecting growth and metabolism of future generations(Carone et al., 2010; Chen et al., 2016; Sharma et al., 2016; Skinner et al., 2013). These few examples suggest that parental environment can have a profound impact on subsequent generations. Elucidating mechanisms behind these environmentally triggered epigenetic programs is essential for a complete understanding of the foundational principles upon which biological inheritance is based.

*Drosophila melanogaster* females, when cohabitated with endoparasitoid wasps, shift to prefer ethanol food as an egglaying substrate, where ethanol food protects Drosophila larvae from wasp (Kacsoh, Lynch, Mortimer, & Schlenke, 2013a). *Drosophila suzukii* similarly shifts egglaying preference to food with atropine, giving its progeny protection against wasp(Poyet et al., 2017). Ethanol preference in *D. melanogaster* is linked to a decrease in Neuropeptide F (NPF) in the female brain (Kacsoh et al., 2013a), consistent with previous work on NPF (Shohat-Ophir, Kaun, Azanchi, Mohammed, & Heberlein, 2012), and its mammalian homolog NPY studied in the context of drug addiction (Gonçalves, Martins, Baptista, Ambrósio, & Silva, 2016; Landayan & Wolf, 2015). NPY modulation governs ethanol consumption in rats (Thiele, Marsh, Marie, Bernstein, & Palmiter, 1998) and is implicated in human alcohol abuse disorders(Mayfield et al., 2002; Mottagui-Tabar et al., 2005). This behavioral output is believed to be a consequence of the NPF/NPY role in the rewards pathway, with NPF signaling being inherently rewarding (Desai, Upadhya, Subhedar, & Kokare, 2013; Shao et al., 2017). NPF activity is considered representative of the motivational state of the fly (Krashes et al., 2009; Landayan & Wolf, 2015). Several recent studies also have shown that ‘stressful’ experiences regulate NPY/NPF levels, providing a link between environmental cues and NPF/NPY signaling (Broqua, Wettstein, Rocher, Gauthier-Martin, & Junien, 1995; Sah et al., 2009; Shohat-Ophir et al., 2012). Here we present findings that link maternal environmental conditions to cause inheritance of an altered reward pathway *via* depressed NPF signaling and preference for ethanol.

## Results

### Inheritance of ethanol preference

Drosophila were cohabitated with female wasps for four days, then separated and flies were placed into embryo collection chambers for 24 hours. Embryos were divided into two cohorts and each developed in the absence of adult flies or wasp. One cohort was used to propagate the next generation and never treated to ethanol food; the second cohort was used in the ethanol preference assay and then discarded (Fig. 1a).

**Figure 1.**
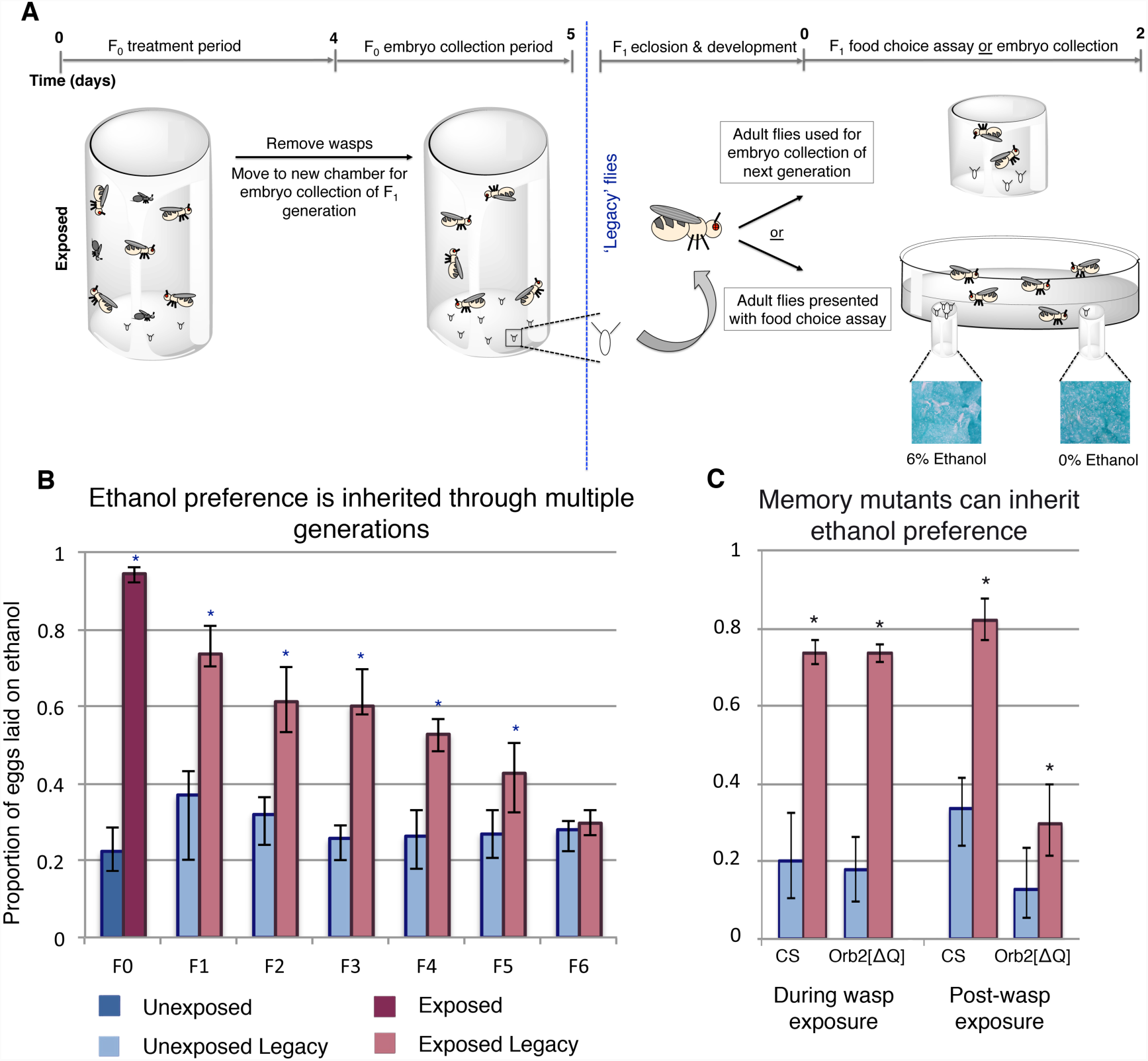
Maternally inherited ethanol preference persists for multiple generations. Schematic of experimental design is shown (**A**). Flies are exposed to wasps for a period of four days prior to egg collection. The descendants from either wasp-exposed or unexposed treatment groups, termed ‘legacy’ flies, are reared until maturity in the absence of both wasps and parental exposure. Legacy flies are either used to propagate the next generation, or are assayed for ethanol preference. Flies from a particular generation are referred to as F_n_, where n denotes the number of generations removed from the treatment. For example, the treatment group itself is F_0_, whereas their direct offspring are F_1_. Ethanol preference is quantified as proportion of eggs laid on ethanol food (**B**), illustrating that this behavior is heritable through the F_5_ generation. Flies with deficient long-term memory were tested for transgenerational inheritance of ethanol (**C**). Asterisk indicates p-value of <0.05.

Wasp-exposed Canton-S flies lay approximately 94% of their eggs on ethanol food (Fig. 1b). This behavior persists in their offspring despite the F_1_ generation never having direct interaction with wasps (Fig. 1b). Ethanol preference in F_1_ was less potent, with 73% of the eggs laid on ethanol food (p = 8.6e^-7^, Table S1). Remarkably, this inherited ethanol preference persisted for five generations, gradually reverting back to the mock exposed baseline (Fig. 1b). These observations were replicated in an additional wild type Oregon R (OR) strain (Table S2), suggesting that the phenomenon is not specific to a particular genetic aberration or background. This indicates that inheritance of ethanol preference is not a permanent germline change, but rather it is a reversible trait. Ethanol preference was measured for two days for initial experiments (Fig. 1-3 & S1); day one and day two showed similar trends, suggesting that flies do not habituate to ethanol nor does the preference fade over the course of the experiment (Table S3).

Confirming previous findings, following a wasp exposure F_0_ flies have an ethanol preference that persists for more than a week, returning to baseline after ten days (Fig. S1a). Sister cohorts of F_1_ flies were collected at two time points along this F_0_ ethanol preference decay; one immediately following wasp exposure (brood 1), and the second ten days post wasp exposure (brood 2). Brood 2 did not display an inherited ethanol preference, suggesting that wasp exposure does not inflict a permanent change in the F_0_ germline (Fig. S1d). Again, these findings replicated in OR flies (Table S2), indicating that these observations are robust and not dependent on the context of a particular genetic background.

To explore further the role of time and dynamics of wasp exposure, multiple generations of flies were exposed to wasps. We found that inherited ethanol preference can be enhanced with successive generations of wasp exposure (Fig. S1e). This trend did not repeat when nonconsecutive generations were repeatedly exposed to wasps (Fig. S1f). This suggests that the enhancing effect observed in the successive exposures is time sensitive and may be linked to the ethanol preference of the parental flies.

To explore the required neural signaling to the germline, mutants defective for long-term memory were assayed. Previous studies have shown that flies defective in long-term memory exhibit an ethanol preference only in the presence of wasps but not after wasp removal. The long-term memory mutant 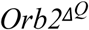 produced offspring with an ethanol preference when the embryo collection was conducted in the presence of wasps, but this ethanol preference was greatly reduced in offspring collected post-wasp exposure (Fig. 1c). These data provide insight in two ways. First, functional long-term memory is not a compulsory requirement to generate ethanol-preferring offspring. Secondly, intact long-term memory is not required to inherit ethanol preference. Given that ethanol preference in the absence of wasps is long-term memory dependent, this experiment reveals that the neuronal signaling is different for maintained ethanol preference in the F_0_ and F_1_ flies(Bozler et al., 2017).

Several other factors point to distinctions between the F_0_ and F_1_ ethanol preference behavior. Male F_1_ legacy flies, mated to naïve females produced offspring with an ethanol preference (Fig. S2a). Additionally, 14-16 day old F_1_ flies displayed an ethanol preference, demonstrating that F_1_ flies do not have an ethanol preference decay curve similar to that of the F_0_ (Fig. S2b).

### Transcriptional changes

Global transcriptional changes in the female head across generations were examined with RNA sequencing. Heads from the F_1_ and F_2_ generation were collected and compared with the F_0_ generation, which was previously reported (Bozler et al., 2017). Analysis of the F_0_ data detected 98 differentially expressed transcripts (15 down and 83 up) (Fig. S3, Table S4). F_1_ and F_2_ heads showed very few differentially expressed transcripts, 4 and 5 transcripts respectively. Of the differentially expressed transcripts, no transcript was shared between groups. These data indicate that although wasp exposure itself results in global transcriptional changes in the female head, this observation does not hold true for the subsequent generations.

### Germline caspases are necessary

Mid-oogenesis germline apoptosis (stage 7-8 oocytes) is triggered upon wasp exposure (Fig. 2a) (Kacsoh, Bozler, & Bosco, 2018; Kacsoh et al., 2018; Kacsoh, Bozler, Ramaswami, & Bosco, 2015). However, this wasp response is not heritable like the ethanol preference behavior, and F_1_ females do not exhibit germline apoptosis (Fig. 2a). Nevertheless, maternal germline knockdown of effector caspases *Dcp-1* and *drice* produce offspring without an ethanol preference, regardless of parental treatment (Fig. 2c). Although protein-starvation triggers germline apoptosis similar to wasp exposure (Fig. 2b), offspring from mothers with starvation-induced apoptosis do not inherit an ethanol preference (Fig. 2d). This indicates that germline apoptosis in and of itself is not sufficient for inheritance of ethanol preference.

**Figure 2.**
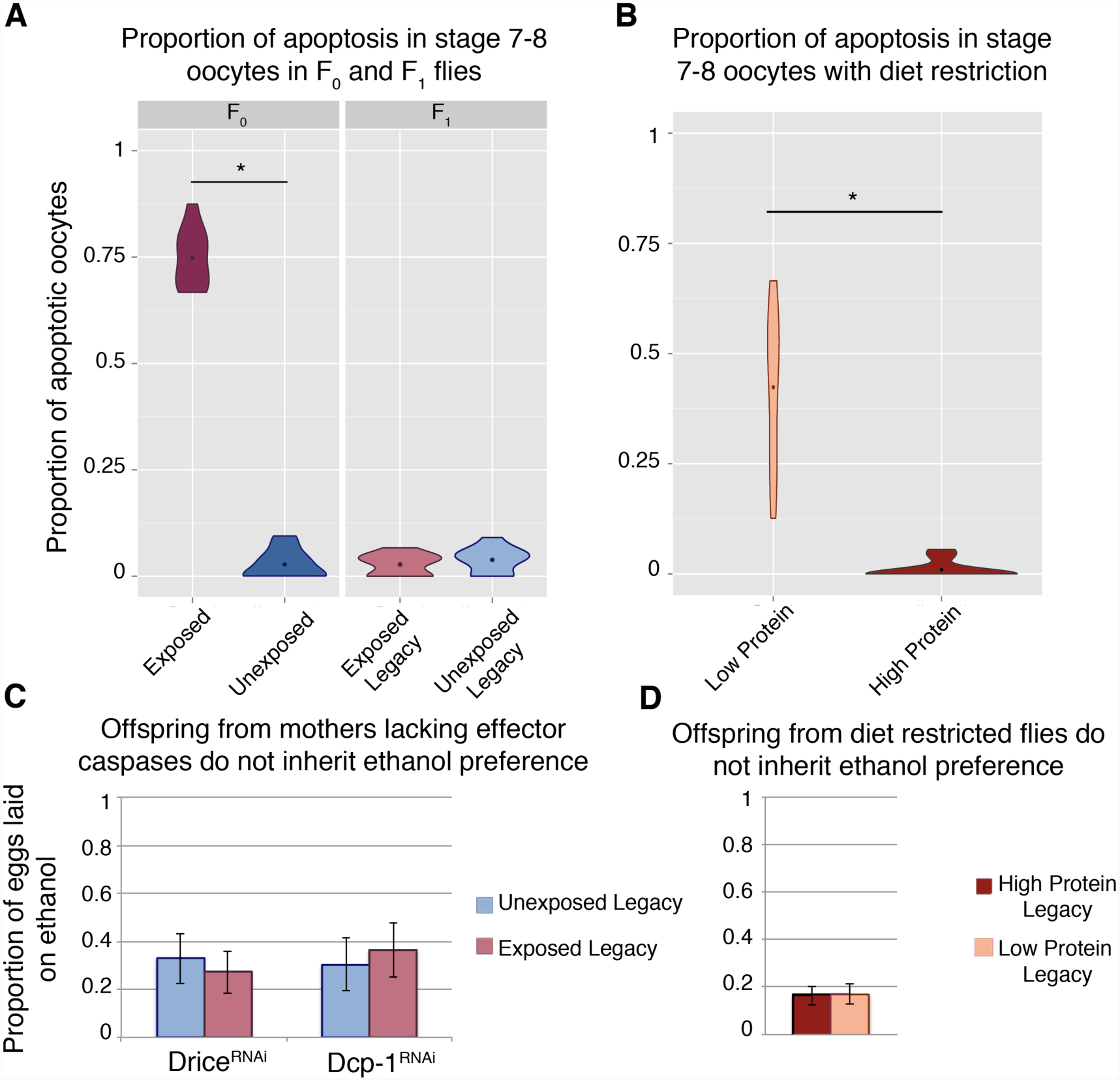
Germline apoptosis and activated caspases play a role in the inheritance of ethanol preference. Apoptosis in stage 7-8 egg chambers was quantified in F_0_ and F_1_ (legacy) flies (**A**). Flies fed a protein-restricted diet have elevated levels of stage 7-8 oocyte apoptosis (**B**). Ethanol preference is not inherited from mothers with Dcp-1 or Drice knockdown (**C**). Offspring from protein-restricted parents don’t inherit an ethanol preference (**D**). Points within violin plots denote the group mean. Asterisk indicates a p-value of <0.05.

### NPF and its receptor modulate germline apoptosis

NPF is known to play a role in food seeking, ethanol consumption, and numerous other reward pathways, and NPF levels decrease in the fan shaped body of female brains following wasp exposure (Kacsoh, Lynch, Mortimer, & Schlenke, 2013b). Even in the presence of wasp overexpression of NPF inhibits ethanol preference, while in the absence of wasp knockdown of NPF is sufficient to induce the ethanol preference behavior (Fig. 3a). Given this NPF modulation of ethanol preference in females we asked whether NPF also signaled to germline cells, triggering caspases and apoptosis. Strikingly, NPF knockdown induces mid-oogenesis apoptosis in the absence of wasps (Fig. 3b), while overexpression of NPF results in no elevation in germline apoptosis even in the presence of wasps (Fig. 3b). Similarly, NPF-receptor (NPFR) knockdown alone leads to significantly elevated levels of apoptosis (28%, when compared to parental line controls p = 6.2e^-4^ & 1.5e^-4^), and this effect is enhanced with wasp exposure (61%, p = 7.0e^-4^) (Fig. 3c). Taken together these observations link ethanol preference behavior and mid-oogenesis apoptosis in the F_0_ females, both processes likely caused by changes in NPF and NPFR signaling.

**Figure 3.**
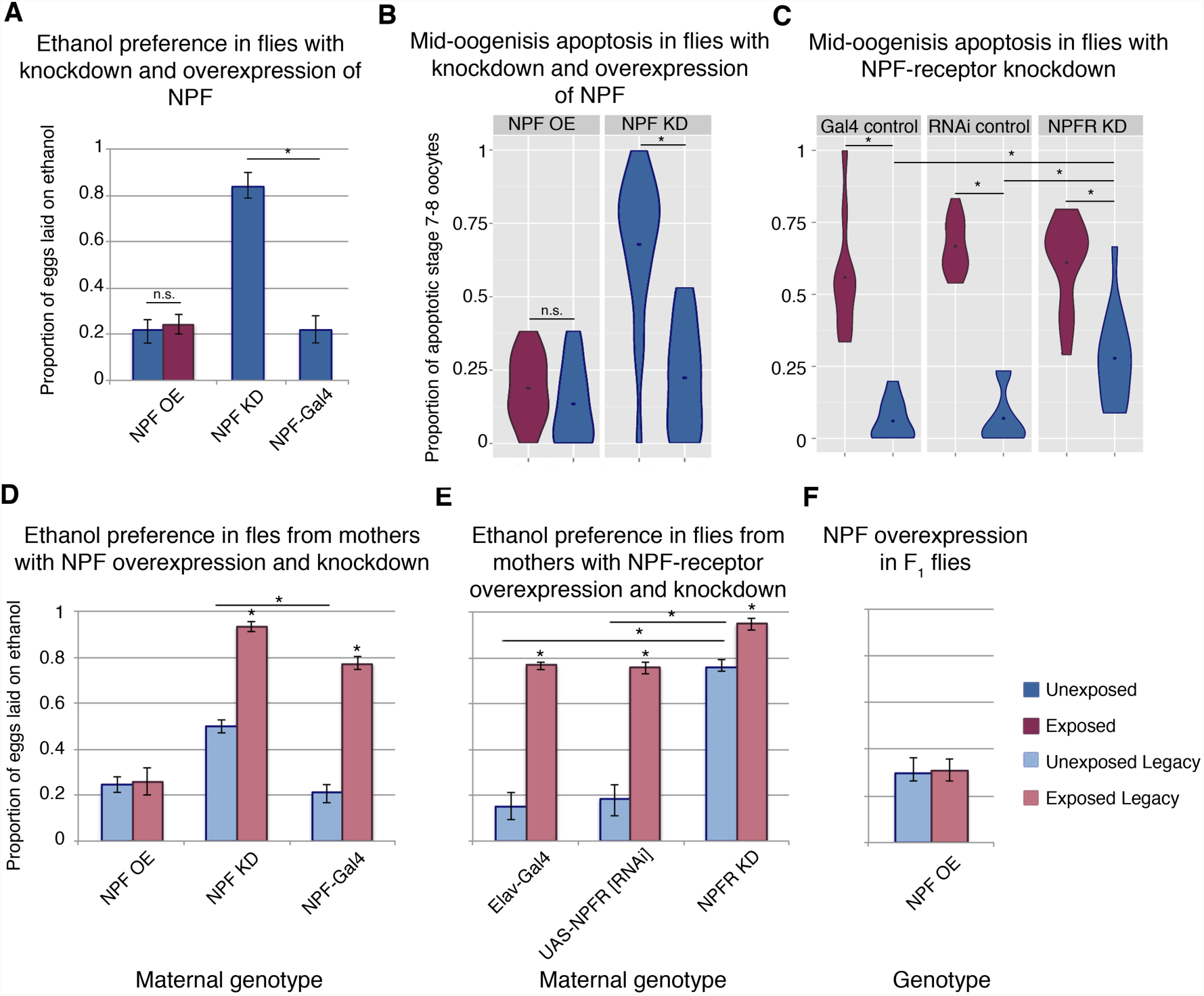
NPF affects ethanol preference and germline apoptosis. NPF overexpression (OE) or knockdown (KD) can alter ethanol preference (**A**). Genetic manipulation of NPF levels can alter levels of germline apoptosis (**B**). Knockdown of NPF receptor leads to increased germline apoptosis (**C**). F_1_ legacy flies have altered ethanol preference depending on the maternal NPF genotype (**D**). F_1_ exposed and unexposed legacy flies inherit ethanol preference from mothers with NPF receptor knockdown (**E**). F_1_ legacy flies overexpressing NPF do not inherit an ethanol preference (**F**). Points within violin plots denote the group mean. Asterisk indicates a p-value of <0.05.

### Changes in NPF trigger transgenerational inheritance of ethanol preference

The NPF-triggered changes in F_0_ behavior and germline also correlate with observed changes in offspring. F_1_ flies from mothers with NPF knockdown exhibit ethanol preference, even in the absence of wasp exposure (Fig. 3d). Inherited ethanol preference is enhanced when the parental NPF knockdown flies are exposed to wasps (Fig. 3d). By contrast, NPF overexpression in F_0_ mothers exposed to wasp produced offspring lacking the ethanol preference (Fig. 3d). NPFR knockdown experiments mirror these findings: Maternal NPFR knockdown produces offspring with an ethanol preference compared to unexposed control lines, again this effect is enhanced when NPFR knockdown is paired with wasp exposure (Fig. 3e). Interestingly, overexpression of NPF in F_1_ flies blocks ethanol preference in the exposed F_1_ legacy group (Fig. 3f), raising the possibility that F_1_ legacy flies inherit NPF in a repressed or low expression state.

We therefore speculated that regulation or depression of NPF might be a means of this behavioral inheritance. Global changes in NPF RNA were not detected in either the F_0_ or F_1_ female heads (Fig. S4). However, antibody staining allowed for a region specific examination of NPF protein levels (Fig. 4a). Anti-NPF signal has clear overlap with the NPF-Gal4 driving the cd8-GFP reporter (Fig. 4a). The fan shaped body has previously been implicated in ethanol preference, and therefore was a focus in this experiment(Kacsoh et al., 2013a). NPF protein levels measured through immunofluorescence were significantly reduced in the fan shaped body of F_0_, F_1_, and F_2_ (two-generations exposed) flies (Fig. 4b). We note that NPF was not reduced in all regions of the F_1_ and F_2_ brains, as intensity of P1 neurons was not reduced in either the F_1_ or F_2_ flies, although significant reduction was observed in P1 neurons of F_0_ flies (Fig. 4c).

**Figure 4.**
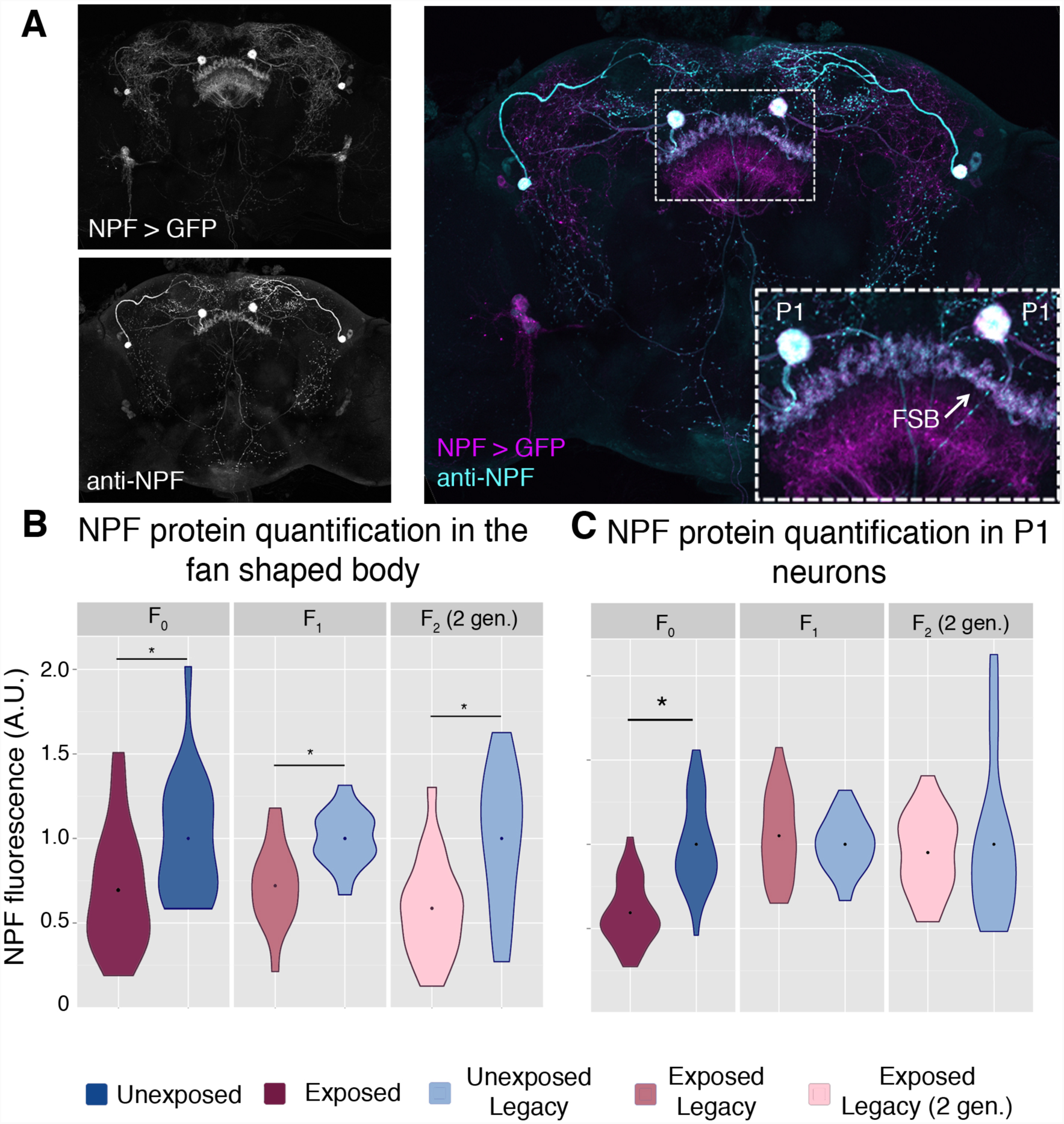
NPF protein is reduced in the fan shaped body following wasp exposure. NPF antibody staining has a similar pattern to that of NPF-Gal4 expression in an adult female brain, inset shows a magnification of the two large P1 neurons and the fan shaped body (FSB)(**A**). NPF protein levels are reduced in the fan shaped body across generations (**B**). NPF depression in P1 neurons is observed only in the F0 generation (**C**). Points within violin plots denote the group mean. Asterisk indicates a p-value of <0.05.

Given the observed link between depressed NPF and oocyte apoptosis, it is notable that F_1_ flies do not have germline apoptosis. It is possible that apoptosis is due to a localized decrease in NPF not shared between the two generations; perhaps the apoptosis is triggered by other NPF neurons or synapses. It is also conceivable that other neural processes are altered in the flies that we did not detect, decoupling the apoptosis and ethanol preference behaviors in the later generations.

### Maternal Chromosomal Inheritance of Ethanol Preference Behavior

To determine whether maternal or paternal exposure were equally important for transgenerational inheritance of ethanol preference, wasp-exposure and mating were controlled in two separate experiments. First, mated females were exposed to wasps in the absence of male flies. Second, wasp-exposed males were mated to naïve virgin females, removing the maternal exposure as a factor. Interestingly, F_1_ offspring from exclusively maternal wasp exposure inherit ethanol preference while F_1_ offspring from exclusively paternal exposures did not (Fig. 5a). We have previously reported that female flies require sight to induce a behavioral response to wasp exposure(Kacsoh et al., 2015). In further support of the maternal contribution to the inheritance of ethanol preference, blind female flies did not produce offspring with an ethanol preference (Fig. 5b). In the reciprocal experiment, blind fathers did generate ethanol-preferring offspring following a wasp exposure (Fig. 5b).

**Figure 5.**
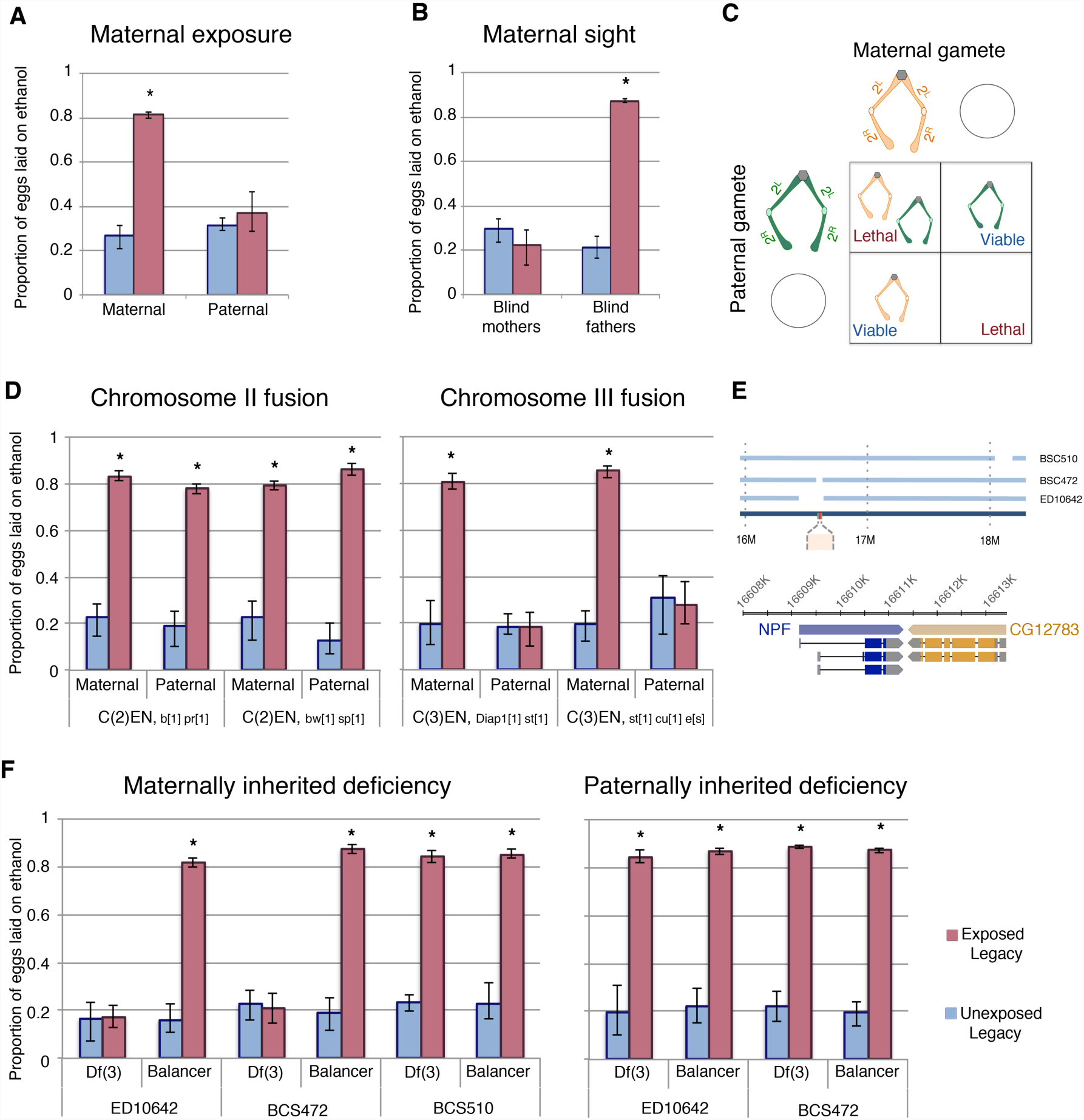
Maternal chromosome 3 is required for inherited ethanol preference. Experiments with exclusively maternal or paternal wasp exposure demonstrate that maternal wasp exposure is necessary for ethanol preference inheritance (**A**). Maternal sight is required for ethanol preference inheritance, but paternal sight is dispensable (**B**). Schematic of compound chromosome 2; progeny inherit both copies from either maternal or paternal source (**C**). Flies receiving either maternal or paternal copies of chromosome 2 have inheritance of ethanol preference, but compound chromosome 3 must be maternally derived to facilitate inheritance of ethanol preference (**D**). Diagram shows the relative location of NPF (red) on chromosome 3 and the deleted region of the deficiency stock (**E**). Inheritance of ethanol preference was observed in flies receiving an intact maternal NPF locus on a balancer chromosome and not in flies from receiving a maternal NPF deficiency (Df3) chromosome: Paternal inheritance of the NPF deficiency had no effect on transmission of ethanol preference (**F**). Asterisk indicates a p-value of <0.05.

**Figure 6.**
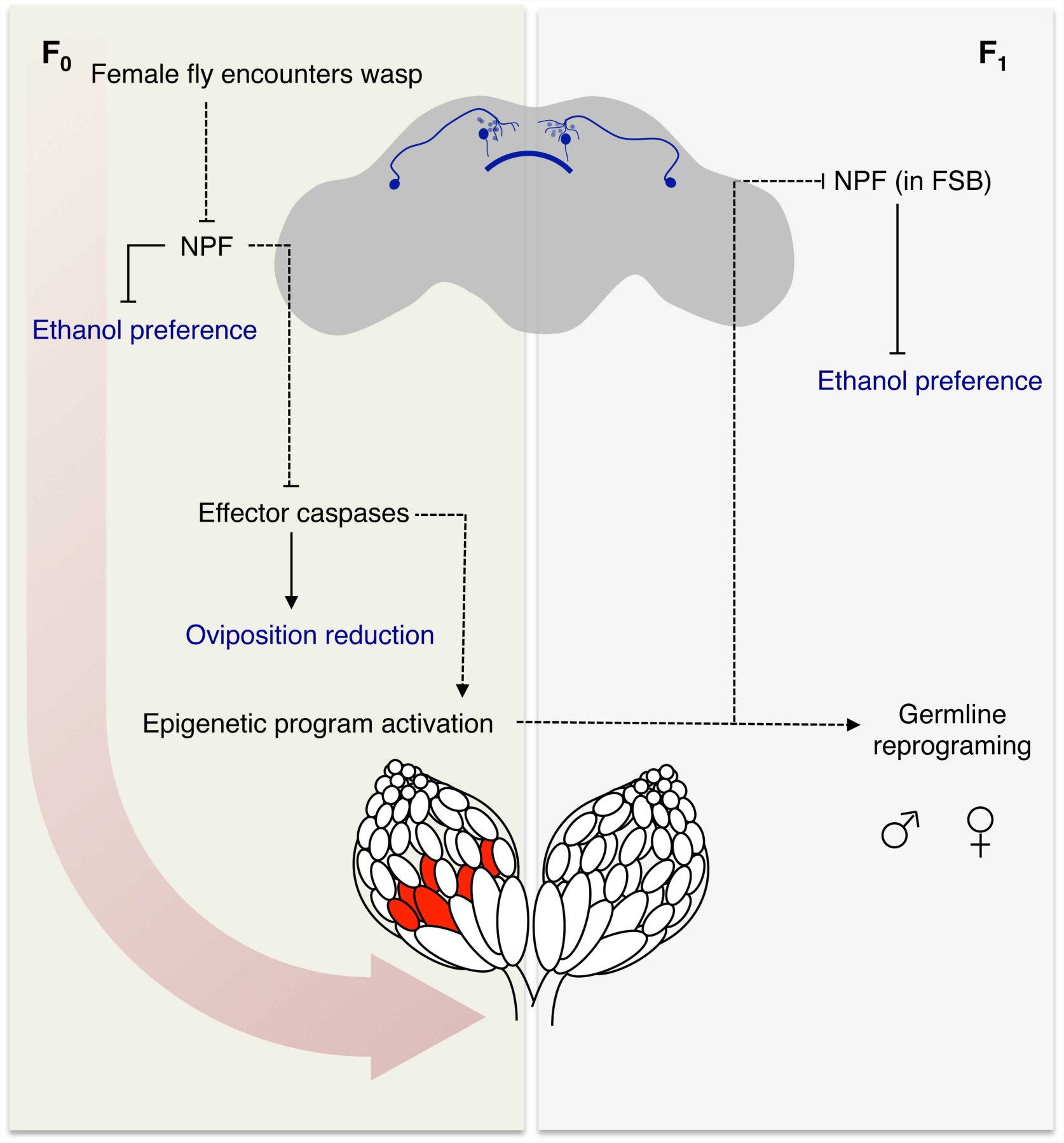
Model for fly-wasp mediated ethanol preference. Wasp encounter leads to a depression of NPF in the female fly brain. Normally NPF inhibits ethanol preference and caspase mediated germline apoptosis. The reduction of NPF triggers ethanol preference and germline caspases. Legacy F_1_ female flies inherit depressed NPF in the FSB, both male and female progeny have altered germline. Measured behavioral outputs are in blue. Dashed lines indicate a speculative or unknown mechanism of action.

Maternal epigenetic inheritance of ethanol preference could be conferred by chromosomal elements and/or cytoplasmic factors. If ethanol preference is inherited through a chromatin mark then chromosome parental-origin tests should reveal a requirement for maternal chromosomal inheritance; however, if inheritance is conferred through cytoplasmic factors, such as noncoding-RNAs, then passage of all chromosomes through paternal gametes should have no effect since wasp-exposed females can still maternally deposit molecules and organelles into the oocyte. To test what maternal components may be conferring inheritance of ethanol preference we first focused on chromosomal elements using attached or compound chromosomes. Flies where each of the two homologs are fused cannot make haplo-chromosome gametes. Instead, they can only make gametes with one or zero copies of the fused chromosome, and therefore F1 flies inherit “pairs” of homologs that are entirely maternally or paternally derived (Fig. 5c). In this manner, we tested each of the two major autosomes for parent-of-origin effects. Using phenotypic markers, flies were sorted as having either a maternal or paternal exclusive homolog pair and assayed for ethanol preference. Chromosome-II fusion flies had similar results when inheriting exclusively maternal or paternal Chromosome-II elements (Fig. 5c). Chromosome-III fusion flies also had inheritance of ethanol preference when receiving both copies of Chromosome-III maternally. However, flies with both copies of Chromosome-III from their fathers failed to inherit an ethanol preference (Fig. 5d). This observation has at least three implications: Most importantly, this indicates that some element on Chromosome-III must be inherited from wasp-exposed mothers in order for ethanol preference behavior to be passed on to F_1_ legacy flies. This also suggests that maternal copies of the Chromosome-X, Chromosome-II or cytoplasmic factors, if important, are not sufficient for inheritance of ethanol preference. Lastly, that oocytes giving rise to eggs with zero copies of maternal Chromosome-II still confer ethanol preferences indicates that exclusion of maternal chromosomes itself does not generally interfere with transgenerational inheritance.

### A Maternal NPF Locus is Required for Epigenetic Inheritance

To further delineate what parts of maternally derived Chromosome-III were required for transgenerational inheritance of ethanol preference we tested chromosomes with well defined deletions. As NPF has previously been shown to control ethanol preference behavior, we speculated that the NPF locus on Chromosome-III may be a target of maternal epigenetic reprogramming(Shohat-Ophir et al., 2012). We also observed that F_1_ legacy flies inherit low levels of NPF expression specifically in the fan shaped body of the brain (Fig. 4a-b), consistent with the possibility that F_1_ flies inherit repressed NPF expression. If the critical maternal Chromosome-III element is the NPF gene locus, then F_1_ offspring with maternal deletions of this chromosomal region may prevent inheritance of ethanol preference, much like not having inherited any maternal copies of Chromosome-III (Fig. 5d). Using females with one Chromosome-III carrying a large deletion of the NPF gene region and one copy of wild-type NPF on a balancer Chromosome-III allowed us to ask whether an intact maternal NPF gene region was necessary for F_1_ inheritance of ethanol preference. We found that legacy F_1_ flies from unexposed mothers had no preference for ethanol, regardless of whether they inherited an intact NPF gene on a balancer chromosome or a chromosomal deletion of the NPF region (Fig. 5e-f). Legacy F_1_ flies from exposed mothers inheriting a wild-type NPF on a balancer chromosome exhibited a strong preference for ethanol, suggesting that multiple rearrangements, deletions and mutations of a balancer Chromosome-III are not sufficient to prevent ethanol preference in F_1_ flies. By contrast, legacy F_1_ flies from exposed mothers inheriting a Chromosome-III deletion of the NPF gene region do not inherit any preference for ethanol (Fig. 5f). This was true for two different Chromosome-III deletions at the NPF locus, whereas a Chromosome-III deletion that does not disrupt the NPF gene had no effect (Fig. 5f). Paternally inherited Chromosome-III deletions were not sufficient to prevent ethanol preference in F_1_ flies (Fig. 5f).

## Discussion

Perhaps the blank slate has more written on it than we once thought. Indeed it would appear that animals are bound to their ancestors in a way that some might consider Lamarckian (Galloway & Etterson, 2007; Herman & Sultan, 2011; J Marshall & Uller, 2007). The ethanol preference we observed in this study is heritable but modifiable and responsive to environmental cues, as it can be enhanced or decay across generations. Our data suggest that there is an ultimate return to pre-wasp exposed state by the F_6_ generation. If there are lingering effects of wasp exposure beyond this generation, they are not detected in our assays. Not only does the ethanol preference behavior revert to unexposed levels, but we also detected no priming or enhancement effect in the F_8_ generation following a second wasp exposure (Fig S1f).

Inheritance of ethanol preference requires several factors: We found that the initiation of the epigenetic program in the founding generation (F_0_) is maternal in nature, and requires effector caspases in the female germline. However, continuation of the epigenetic program throughout the remaining generations is distinctly different in several ways. Both male and female progeny (F_1_) are able to pass on ethanol preference to their offspring. Although, it is possible that the F_1_ generation requires germline effector caspases for the transmission of the ethanol preference, the lack of female germline apoptosis and paternal ability to confer this behavior points to a caspase-independent maintenance mechanism. A further and curious distinction between the generations is in the ethanol preference itself, as it persists in the F_1_ generation, rather than mirroring the F_0_ generation and decaying over 10 days.

The unifying mechanism behind many of these observations is the central role of NPF signaling in this system. Governing both germline apoptosis and the ethanol preference neuronal NPF signaling modulates the ethanol preference as well as its inheritance. Maternal imprinting of the NPF locus or nearby regions has a dominant effect, leading to the possibility that the F_1_ paternal locus is imprinted in *trans*. It is tempting to speculate on the role of canonical imprinting mechanisms, such as the Polycomb repressive complexes, although a molecular apparatus remains elusive for the time being.

This multi-generational ethanol preference underscores the importance of environmental conditions on behavior and physiology. Numerous studies have indicated that we may need to look beyond the individual, to longer lasting and persistent effects of environmental stresses. This study illustrates the complexity of inheritance and highlights the incredible resiliency and plasticity of organisms to adapt to changing circumstance. Of particular interest is the conserved functions of NPF and its mammalian homolog NPY in modulating a variety of human behaviors, including stress responses and alcohol abuse disorders (Thorsell & Mathé, 2017). Our studies raise the intriguing possibility that NPF/NPY and their receptors could be subjected to epigenetically modified states determined by parental environment and experience. Germline inheritance of epigenetically modified neuro-signaling networks, such as those modulated by NPF/NPY, could be one mechanism through which trans-generational inheritance of behavioral predispositions persist, as reported here for Drosophila. It should be noted that such epigenetically inherited behaviors that persist for multiple generations could be interpreted as dominant familial genetic traits. If mammalian NPY is inherited in epigenetically modified states, then this would require a fundamental change in how we study and view inheritance of NPY-related behavioral disorders and possible effects of parental environment.

## Supplementary Materials

### Materials and Methods

#### Fly husbandry

Flies were maintained at room temperature on standard cornmeal-molasses media. A list of fly lines and genotypes used is reported in Table S5. Female flies were considered mature adults at three to five days post eclosion. Flies outside of this age range were not used for experimentation unless specifically noted, as for example in S Fig. 1d. Experiments involving manipulation of the maternal genotype, such as the maternal NPF knockdown, had a crossing scheme to avoid transgene expression in the F_1_ generation. Virgin females with the genotype of interest were crossed to *y,w* males and offspring were scored by eye color to ensure that flies assayed were not carrying both the Gal4 and UAS constructs.

#### Wasp-exposure

Mature adult flies were used for wasp exposures: 40 female flies, 10 male flies, and 20 female Lh14 (*Leptopilina heterotoma*) wasps were placed in a vial with cornmeal-molasses media. This cohabitation (wasp exposure period) lasted for four days. The unexposed control consisted of the 40 female flies and 10 male flies with no wasp cohabitation. Both treatment groups were maintained at room temperature (approximately 22° C) with a 12 hour light-dark cycle for the duration of the exposure period.

At the conclusion of the exposure period, flies were separated into two cohorts. Following the removal of all wasps, one group of flies was used to propagate the next generation, while the second group was assayed for ethanol preference. Group one was placed on molasses-based embryo collection plates, supplemented with yeast paste, for egg collection. The collection period lasted for 24-hours, at which point the adult flies were removed. First instar larvae were transferred from these embryo plates to standard media vials. Larvae were density controlled to approximately 40 larvae per vial.

The second group was assayed for ethanol preference using a food-choice assay (Kacsoh, Bozler, Hodge, Ramaswami, & Bosco, 2015). Briefly, five female flies and one male fly were placed into a modified petri dish with mesh top, termed the ‘fly corral’. Two food sources were placed at opposite ends of the ‘fly corral’. Each food source consisted of 0.45 g of instant drosophila media, hydrated with 2mL liquid. Control food was hydrated entirely with distilled water, where as ethanol food was prepared with distilled water and a final addition of 95% ethanol to the top of the prepared food, creating a food with 6% ethanol by volume. Food sources were removed and replaced after 24 hours. Figures report the egg laying behavior of the first 24-hour interval unless otherwise noted. Total number of eggs laid on each food source was counted in a blinded fashion with treatment unknown to the counter. These egg counts are reported as a proportion of eggs laid on ethanol food. Flies that encountered ethanol-containing food were excluded from additional experimentation or lineage propagation. Fly corral experiments had ten replicates (cages) per condition.

#### Transgenerational behavior experiments

Legacy flies, those descending from either the unexposed or exposed treatment, were divided into cohorts as described above for behavioral assay or embryo collection. These flies were not re-exposed to wasps except in the instance of multigenerational exposure experiments. Two experiments were conducted that involved multiple generations of treatment. For the successive exposures, three groups of flies were assayed; exposed legacy (2 generations), exposed legacy (1 generation), and unexposed legacy. In this instance, the exposed legacy (2 generations) group was generated by subjecting F_1_ exposed legacy flies to an additional round of wasp exposures. These flies therefore had grandparental and parental wasp exposure. Exposed legacy (1 generation) had parental wasp exposure only (Figure 3 B). It is important to note that the parents of the ‘exposed legacy (1 generation)’ flies were F_1_ unexposed legacy flies, and therefore had the same density control and egg collection as the other groups for the multigenerational duration of the experiment.

It is critical to note that baseline ethanol preference is highly variable depending on environmental conditions. Key factors are temperature and humidity, all ethanol oviposition assays were conducted in an environmentally controlled room at 25°C, approximately 30% humidity (+/- 10%) with overhead lighting and a 12-hour light/dark cycle. Despite these controls, baseline ethanol preference varies day-to-day. For this reason, all groups for direct comparison (used in statistical tests) were tested at the same time.

Pertaining to the nonconsecutive exposure experiments; again three groups were assayed, the exposed legacy F_8_ (2 generations), exposed legacy (1 generation), and the unexposed legacy. For these experiments, the exposed legacy F_8_ (2 generations) group was created by subjecting F_7_-exposed legacy flies to an additional round of wasp exposures. These flies had a six-generation gap between ancestral wasp exposures. Flies in the exposed legacy (1 generation) group were produced by exposing F_7_ unexposed legacy flies to wasps, and collecting the subsequent offspring.

Several experiment specific modifications were made to the methods described above. To parse the maternal and paternal contributions to the inheritance of ethanol preference two experiments were conducted. First, 40 mated female flies were used for wasp exposure, in the absence of males. Ten males were added to the population for the embryo collection period. For paternal contribution, male flies were removed from the exposure chamber and mated to unexposed virgin females. To test the role of vision in maternal inheritance, blind female flies mutant in ninaB, were crossed to wild type (CS) males. The reciprocal experiment crossed ninaB[1] males to CS female. These experiments were run in parallel and wasp exposures were preformed as previously described.

Compound chromosome experiments crossed two fusion stocks together (either chromosome-II or chromosome-III). The fusion lines retained phenotypic markers, and offspring with maternal or paternal chromosomes were sorted accordingly. Deficiency lines were crossed to CS flies and the genotype of the offspring (balancer or deficiency) was inferred from phenotypic markers.

Particular modifications for the 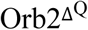 memory-mutant experiments included an extra day of embryo collection. Following three-days of wasp exposure, flies and wasps were moved to the embryo collection chamber for the final treatment day. Eggs were collected for 24-hours in the presence of the 20 female Lh14 wasps. At the end of this period, wasps were removed and a new embryo collection plate was introduced for the second day of embryo collections. This second day of collection corresponds to the standard embryo collection timeframe in the above-described experiments. F_1_ flies had the same genotype as the parental line.

Sibling cohorts were collected to assess the longevity of the germline change. ‘Brood 1’ flies were collected in the 24-hours immediately following the removal of the wasps. ‘Brood 2’ flies were collected from the same parents, 10 days after the termination of the wasp exposure.

Finally, diet restriction experiments had two groups one with high protein and the other low protein diets. Low protein flies were maintained on molasses based embryo plates. The high protein group was maintained in similar fashion, but with the addition of yeast paste. High/low diet was maintained for four days prior to embryo collection.

#### Apoptosis quantification

Following the treatment period, ovaries were dissected and fixed in 4% formaldehyde for 30 minutes. Samples were stained with DAPI and apoptosis was scored based on the morphology of the nurse cell DNA. A researcher blinded to the genotype and treatment group of the samples preformed the scoring. At a minimum, 15 ovaries were scored across 3 replicates (independent wasp exposures) for each group.

#### Immunostaining and microscopy

Antibody to neuropeptide F was generated in a rabbit to the full length NPF peptide: C-Ahx-SNSRPPRKNDVNTMADAYKFLQDLDTYYGDRARVRFamide. The antibody was subsequently purified using a truncated peptide containing the first 28 amino acids of NPF. Following purification, the antibody was depleted using a peptide of the eight amino acid C-terminal tail, shared by many neuropeptides. All peptide synthesis, antigen injection, serum preparation and peptide purification and depletions were performed by 21^st^ Century Biochemicals.

Whole flies were fixed in 4% formaldehyde overnight at 4° C. Female brains were dissected, blocked, and incubated with anti-NPF (1:1000) overnight at 4° C. Antibody solution was removed and samples were blocked before the addition of the secondary antibody, anti-rabbit 488 (1:200), at room temperature for two-hours. Samples were counter stained with DAPI.

For NPF quantification, flies expressing a RFP tagged histone were dissected along with treatment groups and stained in the same solution. Pixel intensity of the fan shaped body (FSB) was measured in Image J. The FSB was outlined by hand and intensity measured. A background measure was made of the region immediately ventral to the FSB, with the same total area as the outlined FSB. The background value was subtracted from FSB measurement. Finally, the background-adjusted intensity value for each brain was divided by the arc length of its’ FSB. This process was repeated for each treatment group and the corresponding histone-RFP flies. These values were normalized to the histone-RFP flies to serve as a control for batch specific variation in staining. Each treatment group was normalized to the unexposed average of that replicate using the formula(s):

Fluorescence = (FSB_intensity_-background_intensity_)/FSB_length_

BatchNormalized=(Fluorescence_CantonS_/Fluorescence[avg]_his-RFP_)

AFU=BatchNormalized_exposed_/BatchNormalized[avg]_unexposed_

Standard fluorescent images were visualized with the Nikon Eclipse E800 microscope and the Olympus DP71 camera. For each experiment, wasp exposure and staining were performed on two separate occasions and final data was pooled after checking for the absence of a batch effect. A minimum of 10 brains were dissected for each treatment replicate as well as RFP-histone co-staining brains. Final quantified sample size range from 15 to 20 (normalized brains), due to sample loss or damage. Imaged samples were only excluded if clear damage or trauma (from dissection or staining process) was evident in the region of interest (FSB or P1 nuerons).

#### RNA quantification

Mature female flies were anesthetized with CO_2_ and collected in 15 mL conical tubes, either immediately following the treatment period (F_0_), or 3-5 days post eclosion (F_1_-F_2_). Flies were frozen in liquid nitrogen and briefly vortexed to separate whole heads. Approximately 100 heads were collected for each replicate. A miRNeasy Kit (Qiagen) with on-column DNase treatment was used for RNA isolation. Four samples of each treatment group were prepared.

RNA samples were depleted of rRNA followed by random priming. Minimum sequencing depth per sample was 40 million paired-end reads on the Illumina platform. Sequencing reads were indexed to transcripts using Kallisto and the Ensembl genome (BDGP6) with 100 bootstraps (Aken et al., 2016; Bray, Pimentel, Melsted, & Pachter, 2016). Downstream processing and statistical analyses used Sleuth (Pimentel, Bray, Puente, Melsted, & Pachter, 2017). Heat maps were generated using hierarchical clustering and the R package pheatmap.

NPF transcript was measured by qPCR (SYBR Green, Thermo-Fisher 4309155). NPF primer targeted mRNA (TCCTGGTTGCCTGTGTGG, TCAGCCATAGTGTTGACATCG). Actin served as the control gene (CGCAAGGATCTGTATGCCAA, ACGGAGTACTTGCGCTCTGG). Fold change was calculated using the delta-delta Ct method.

#### Statistics

Statistical tests were run in R (3.0.2 version, ‘Frisbee Sailing’). P-values for egg count data, NPF staining, and apoptosis quantification, were produced by applying a Mann-Whitney Rank Sum test. Error bars presented in the egg count ethanol preference graphs are bootstrap confidence intervals, generated using the boot package.

---

**Figure S1.**
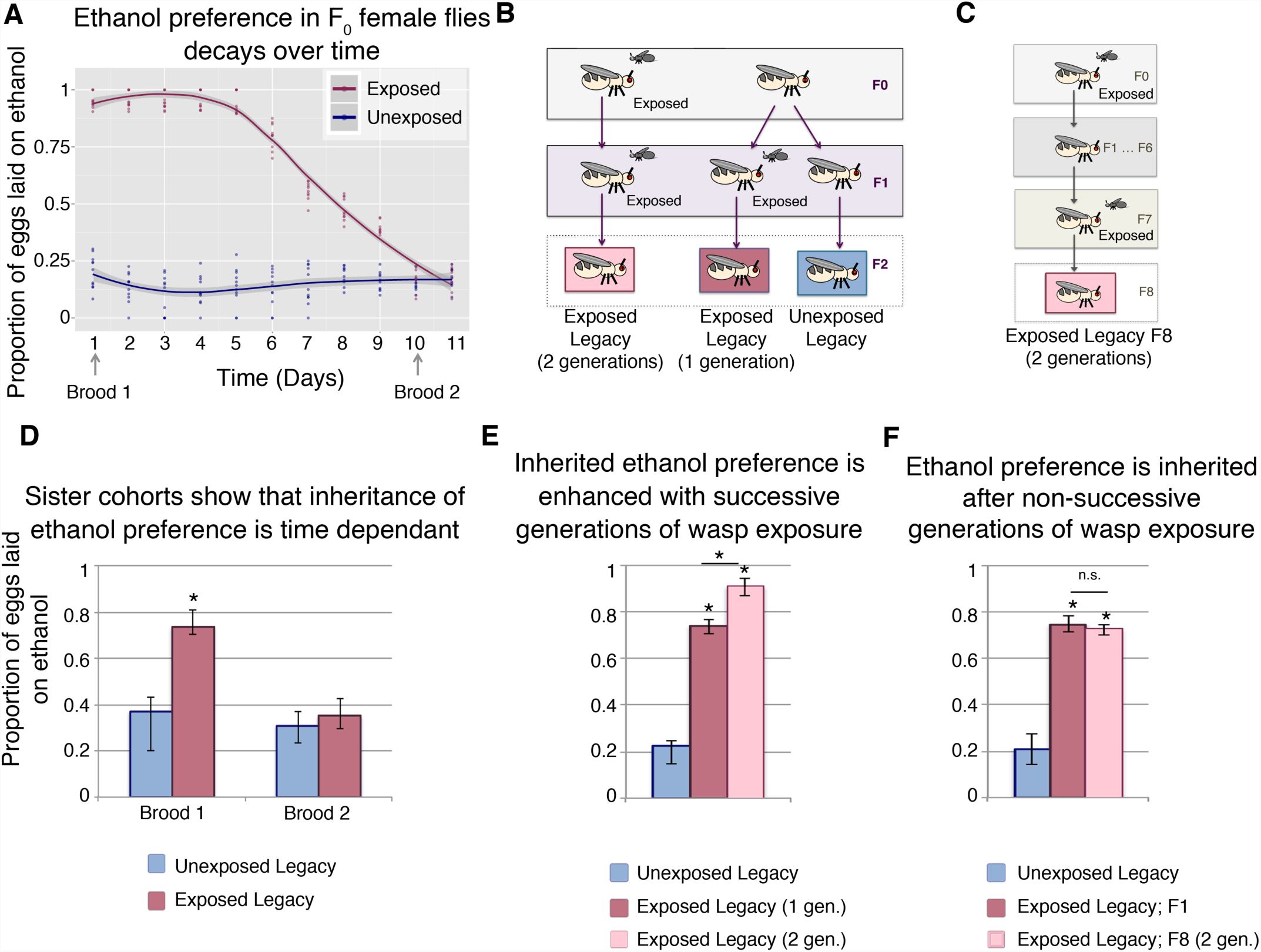
Temporal dynamics of wasp exposure effect inheritance of ethanol preference. Ethanol preference decays following wasp exposure (in F_0_ flies), with loess regression, shaded region indicates standard error (**A**). Diagram of multigenerational exposure is shown for successive generations (**B**), and non-consecutive generations (**C**). Quantification of ethanol preference from sister cohorts collected at different intervals post-wasp exposure (**D**). Flies with successive generations of wasp exposure have enhanced ethanol preference (**E**). Alternatively, flies from a second generation of non-consecutive wasp exposure (exposure of F_7_ flies) exhibit an ethanol preference similar to that of one-generation wasp exposed flies (**F**). Asterisk indicates a p-value of <0.05.

**Figure S2.**
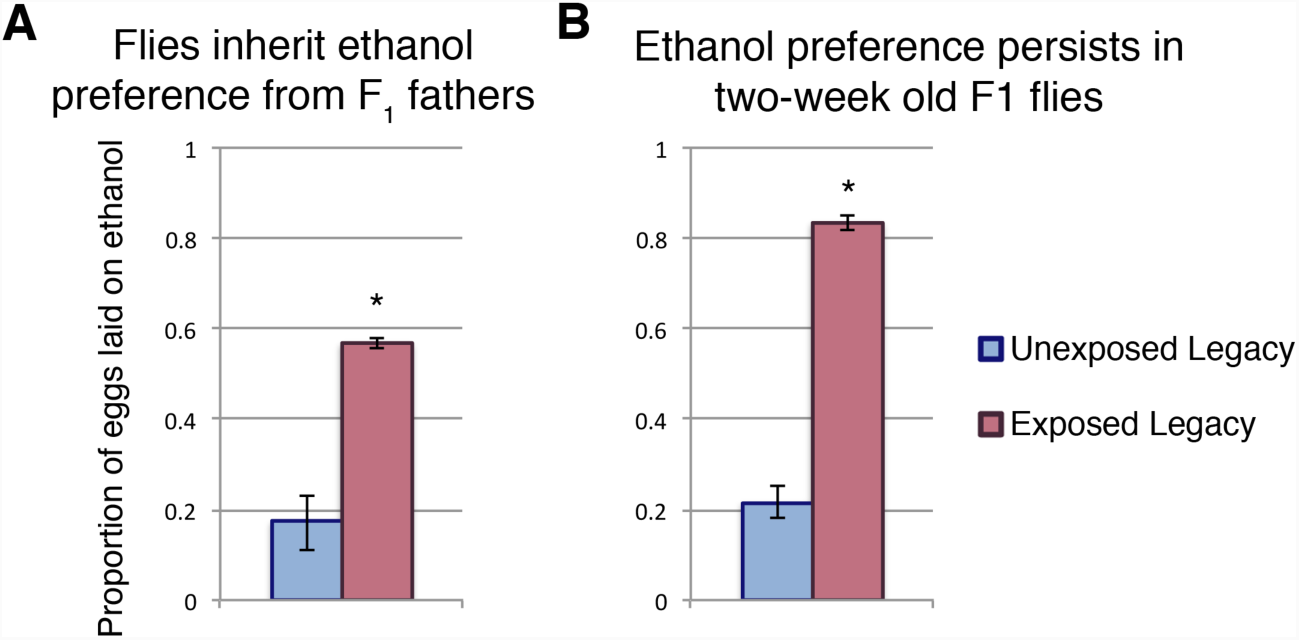
F_1_ ethanol preference has distinct characteristics from those of the parental F_0_ generation. Male F_1_ flies are able to pass on ethanol preference to their offspring (**A**). Ethanol preference of F_1_ flies has not decayed two-weeks post eclosion (**B**). Asterisk indicates a p-value of <0.05.

**Figure S3.**
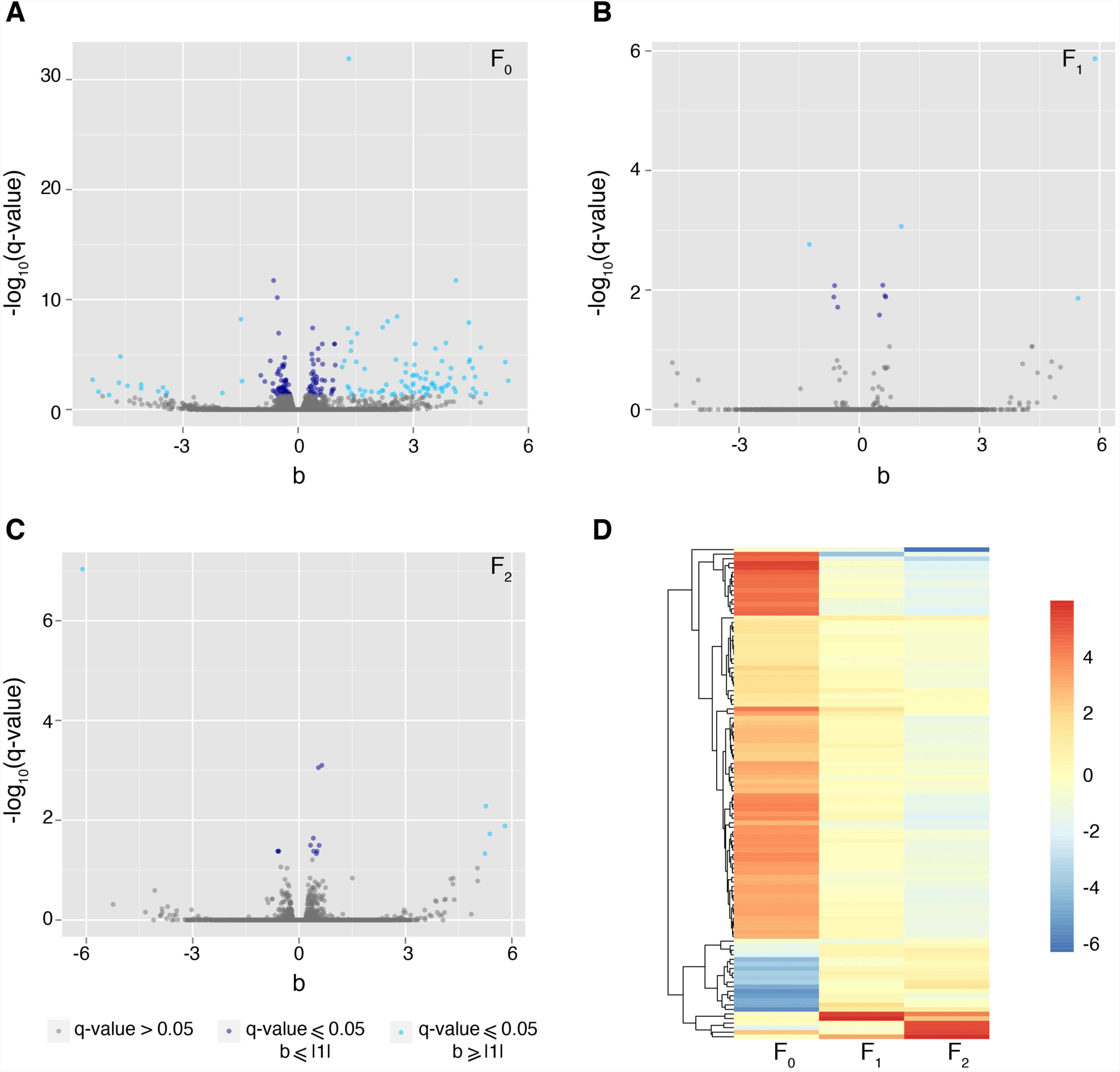
Global transcriptional changes in the female head. RNA sequencing was preformed on heads of F_0_, F_1_, and F_2_ females. Volcano plots show the distribution of transcript expression and significance. F_0_ flies have a considerable number of differentially expressed transcripts (**A**). Where as F_1_ and F_2_ heads have very few changes in transcripts (**B**) & (**C**). The beta value is approximately analogous to the natural log fold change of the transcript, and the q-value is the measure of significance. Grey points indicate a transcript with non-significant q-value, dark blue points indicate transcripts with significant q-value but that do not meet the beta value threshold. Light blue dots have significant q-value and an absolve value of beta greater or equal to one. Heat map shows the trend of transcript expression over the three generations (**D**). Transcript meeting the threshold criteria (q-value and beta) for any one generation was included in the map.

**Figure S4.**
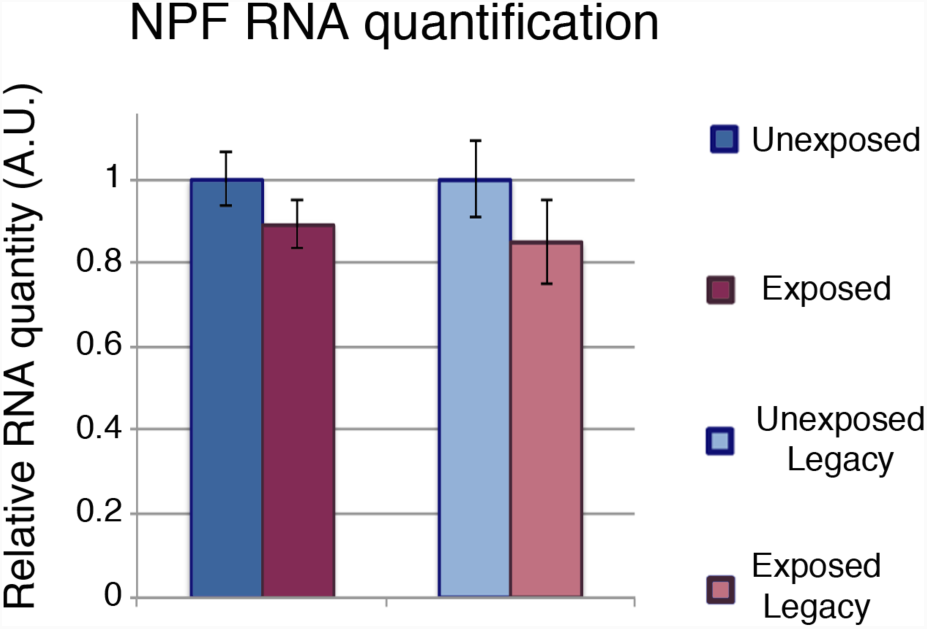
mRNA quantification of NPF in female fly heads. Asterisk indicates a p-value of <0.05.

**Table S1.** Statistical tests and p-values relating to main text figures.

| Figure | Comparison groups | p-value | statistical test |
| --- | --- | --- | --- |
| 1B | F0 (Exposed vs unexposed) | 1.08E-05 | Mann-Whitney Rank Sum |
| 1B | F1 (Exposed vs unexposed) | 8.64E-07 | Mann-Whitney Rank Sum |
| 1B | F2 (Exposed vs unexposed) | 2.58E-08 | Mann-Whitney Rank Sum |
| 1B | F3 (Exposed vs unexposed) | 1.29E-08 | Mann-Whitney Rank Sum |
| 1B | F4 (Exposed vs unexposed) | 4.13E-06 | Mann-Whitney Rank Sum |
| 1B | F5 (Exposed vs unexposed) | 0.00833 | Mann-Whitney Rank Sum |
| 1B | F6 (Exposed vs unexposed) | 0.6063 | Mann-Whitney Rank Sum |
| 1C | CS-during (Exposed vs unexposed) | 0.0001817 | Mann-Whitney Rank Sum |
| 1C | Orb2-during (Exposed vs unexposed) | 0.000278 | Mann-Whitney Rank Sum |
| 1C | CS-post (Exposed vs unexposed) | 1.08E-05 | Mann-Whitney Rank Sum |
| 1C | Orb2-post (Exposed vs unexposed) | 0.02065 | Mann-Whitney Rank Sum |
| 2A | F0 (Exposed vs unexposed) | 0.0001593 | Mann-Whitney Rank Sum |
| 2A | F1 (Exposed vs unexposed) | 0.3144 | Mann-Whitney Rank Sum |
| 2B | High protein vs low protein | 0.0001079 | Mann-Whitney Rank Sum |
| 2C | Drice[RNAi] (Exposed vs unexposed) | 0.5787 | Mann-Whitney Rank Sum |
| 2C | Dcp-1[RNAi] (Exposed vs unexposed) | 0.05889 | Mann-Whitney Rank Sum |
| 2D | High protein vs low protein | 0.933864 | Mann-Whitney Rank Sum |
| 3A | NPF OE (Exposed vs unexposed) | 0.5787 | Mann-Whitney Rank Sum |
| 3A | NPF KD vs NPF-Gal4 | 1.08E-05 | Mann-Whitney Rank Sum |
| 3B | NPF OE (Exposed vs unexposed) | 0.1758 | Mann-Whitney Rank Sum |
| 3B | NPF KD vs NPF-Gal4 | 1.76E-05 | Mann-Whitney Rank Sum |
| 3C | Elav-Gal4 (Exposed vs unexposed) | 0.0001697 | Mann-Whitney Rank Sum |
| 3C | NPFR[RNAi] (Exposed vs unexposed) | 3.07E-06 | Mann-Whitney Rank Sum |
| 3C | NPFR KD (Exposed vs unexposed) | 0.0007069 | Mann-Whitney Rank Sum |
| 3C | NPFR KD vs NPFR[RNAi] unexposed | 0.0001503 | Mann-Whitney Rank Sum |
| 3C | NPFR KD vs Elav-Gal4 unexposed | 0.0006232 | Mann-Whitney Rank Sum |
| 3D | NPF OE (Exposed vs unexposed) | 0.4359 | Mann-Whitney Rank Sum |
| 3D | NPF KD (Exposed vs unexposed) | 1.08E-05 | Mann-Whitney Rank Sum |
| 3D | NPF-Gal4 (Exposed vs unexposed) | 1.08E-05 | Mann-Whitney Rank Sum |
| 3D | NPF KD vs NPF-Gal4 unexposed | 1.08E-05 | Mann-Whitney Rank Sum |
| 3E | Elav-Gal4 (Exposed vs unexposed) | 0.0001817 | Mann-Whitney Rank Sum |
| 3E | NPFR[RNAi] (Exposed vs unexposed) | 0.0001817 | Mann-Whitney Rank Sum |
| 3E | NPFR KD (Exposed vs unexposed) | 0.0001817 | Mann-Whitney Rank Sum |
| 3E | NPFR KD vs NPFR[RNAi] unexposed | 1.08E-05 | Mann-Whitney Rank Sum |
| 3E | NPFR KD vs Elav-Gal4 unexposed | 0.0001817 | Mann-Whitney Rank Sum |
| 3F | NPF OE (Exposed vs unexposed) | 0.7333 | Mann-Whitney Rank Sum |
| 4B | F0 (Exposed vs unexposed) | 0.009027 | Mann-Whitney Rank Sum |
| 4B | F1 (Exposed vs unexposed) | 0.0004949 | Mann-Whitney Rank Sum |
| 4B | F2 (Exposed vs unexposed) | 0.002572 | Mann-Whitney Rank Sum |
| 4C | F0 (Exposed vs unexposed) | 3.09E-11 | Mann-Whitney Rank Sum |
| 4C | F1 (Exposed vs unexposed) | 0.3972 | Mann-Whitney Rank Sum |
| 4C | F2 (Exposed vs unexposed) | 0.6378 | Mann-Whitney Rank Sum |
| 5A | maternal (Exposed vs unexposed) | 0.000011 | Mann-Whitney Rank Sum |
| 5A | paternal (Exposed vs unexposed) | 0.1904 | Mann-Whitney Rank Sum |
| 5B | Blind mothers (Exposed vs unexposed) | 0.3154 | Mann-Whitney Rank Sum |
| 5B | Blind fathers (Exposed vs unexposed) | 0.0002712 | Mann-Whitney Rank Sum |
| 5D | Chr-II Maternal C(2)EN b[1] pr[1] (Exposed vs unexposed) | 0.0001796 | Mann-Whitney Rank Sum |
| 5D | Chr-II Paternal C(2)EN b[1] pr[1] (Exposed vs unexposed) | 0.0002695 | Mann-Whitney Rank Sum |
| 5D | Chr-II Maternal C(2)EN bw[1] sp[1] (Exposed vs unexp | 0.0004456 | Mann-Whitney Rank Sum |
| 5D | Chr-II Paternal C(2)EN bw[1] sp[1] (Exposed vs unexp | 0.0002695 | Mann-Whitney Rank Sum |
| 5D | Chr-III Maternal C(3)EN Diap1[1] sp[1] (Exposed vs unexposed) | 0.0006306 | Mann-Whitney Rank Sum |
| 5D | Chr-III Paternal C(3)EN Diap1[1] sp[1] (Exposed vs unexposed) | 0.7308 | Mann-Whitney Rank Sum |
| 5D | Chr-III Maternal C(3)EN st[1] cu[1] e[s] (Exposed vs unexposed) | 0.0006258 | Mann-Whitney Rank Sum |
| 5D | Chr-III Paternal C(3)EN st[1] cu[1] e[s] (Exposed vs unexposed) | 0.8857 | Mann-Whitney Rank Sum |
| 5F | Maternal Df(3)ED10642 (Exposed vs unexposed) | 0.8796 | Mann-Whitney Rank Sum |
| 5F | Maternal ED10642-balancer (Exposed vs unexposed) | 0.0001766 | Mann-Whitney Rank Sum |
| 5F | Maternal Df(3)BCS472 (Exposed vs unexposed) | 0.7569 | Mann-Whitney Rank Sum |
| 5F | Maternal BCS472-balancer (Exposed vs unexposed) | 0.000278 | Mann-Whitney Rank Sum |
| 5F | Maternal Df(3)BCS510 (Exposed vs unexposed) | 0.002141 | Mann-Whitney Rank Sum |
| 5F | Maternal BCS510-balancer (Exposed vs unexposed) | 0.002141 | Mann-Whitney Rank Sum |
| 5F | Paternal Df(3)ED10642 (Exposed vs unexposed) | 0.0001756 | Mann-Whitney Rank Sum |
| 5F | Paternal ED10642-balancer (Exposed vs unexposed) | 0.0001796 | Mann-Whitney Rank Sum |
| 5F | Paternal Df(3)BCS472 (Exposed vs unexposed) | 0.0004426 | Mann-Whitney Rank Sum |
| 5F | Paternal BCS472-balancer (Exposed vs unexposed) | 0.0002451 | Mann-Whitney Rank Sum |
| S1D | Brood 1 (Exposed vs unexposed) | 2.49E-10 | Mann-Whitney Rank Sum |
| S1D | Brood 2 (Exposed vs unexposed) | 0.6305 | Mann-Whitney Rank Sum |
| S1E | Exposed (1 gen) vs unexposed | 1.08E-05 | Mann-Whitney Rank Sum |
| S1E | Exposed (2 gen) vs unexposed | 1.08E-05 | Mann-Whitney Rank Sum |
| S1E | Exposed (1 gen) vs exposed (2 gen) | 1.08E-05 | Mann-Whitney Rank Sum |
| S1F | Exposed (1 gen) vs unexposed | 1.82E-04 | Mann-Whitney Rank Sum |
| S1F | Exposed F8 (2 gen) vs unexposed | 1.08E-05 | Mann-Whitney Rank Sum |
| S1F | Exposed (1 gen) vs exposed F8 (2 gen) | 0.472 | Mann-Whitney Rank Sum |
| S2A | Exposed vs unexposed | 1.08E-05 | Mann-Whitney Rank Sum |
| S2B | Exposed vs unexposed | 0.000181 | Mann-Whitney Rank Sum |
| S4 | F0 (Exposed vs unexposed) | 0.3429 | Mann-Whitney Rank Sum |
| S4 | F1 (Exposed vs unexposed) | 0.3429 | Mann-Whitney Rank Sum |

**Table S2.** Oregon R experimental data. Key experiments were replicated using the additional wild-type strain OreR. “Corresponding Figure” indicates the experiment that was replicated: A listing of Fig1B therefore indicates that the experimental conditions for Figure 1B were duplicated using OreR flies.

|  |  | Day 1 |  |  | Day 2 |  |  |
| --- | --- | --- | --- | --- | --- | --- | --- |
| Corresponding Figure for Duplicate Experiment | Description | Mean (experimental) | Mean (control) | p-value | Mean (experimental) | Mean (control) | p-value |
| 1B | F0 | 0.913 | 0.288 | 1.08E-05 | 0.927 | 0.276 | 1.08E-05 |
| 1B | F1 | 0.704 | 0.303 | 1.08E-05 | 0.695 | 0.276 | 1.81E-04 |
| 1B | F2 | 0.679 | 0.286 | 1.08E-05 | 0.691 | 0.271 | 1.82E-04 |
| 1B | F3 | 0.675 | 0.25 | 1.08E-05 | 0.647 | 0.266 | 1.82E-04 |
| 1B | F4 | 0.556 | 0.211 | 2.44E-04 | 0.534 | 0.221 | 1.08E-05 |
| 1B | F5 | 0.366 | 0.277 | 0.07526 | 0.41 | 0.221 | 0.000129 |
| 1B | F6 | 0.273 | 0.263 | 0.7394 | 0.224 | 0.238 | 0.6305 |
| Not shown in figure | F7 | 0.202 | 0.208 | 0.4359 | 0.211 | 0.193 | 0.3429 |
| 2A | F0 apoptosis (Exposed vs unexposed) | 0.705 | 1.70E-02 | 0.000144 |  |  |  |
| 2A | F1 apoptosis (Exposed vs unexposed) | 0.017 | 3.10E-02 | 0.2931 |  |  |  |
| 5A | Paternal | 0.36 | 0.45 | 0.1655 | 0.34 | 0.35 | 0.8534 |
| S1D | Brood 2 | 0.23 | 0.27 | 0.705 | 0.24 | 0.27 | 0.6842 |
| S1E | Exposed (1 gen) vs unexposed | 0.744 | 0.213 | 1.81E-04 | 0.723 | 0.186 | 1.82E-04 |
| S1E | Exposed (2 gen) vs unexposed | 0.91 | - | 1.08E-05 | 0.924 | - | 1.82E-04 |
| S1E | Exposed (1 gen) vs exposed (2 gen) | - | - | 1.81E-04 | - | - | 1.81E-04 |
| S1F | Exposed (1 gen) vs unexposed | 0.79 | 0.258 | 1.82E-04 | 0.81 | 0.223 | 1.82E-04 |
| S1F | Exposed F8 (2 gen) vs unexposed | 0.76 | - | 1.82E-04 | 0.792 | - | 1.80E-04 |
| S1F | Exposed (1 gen) vs exposed F8 (2 gen) | - | - | 0.1209 | - | - | 0.7048 |

**Table S3.** Canton S day-2 data; mean(s) and p-value(s).

| Corresponding Figure | Description | Mean (experimental) | Mean (control) | p-value |
| --- | --- | --- | --- | --- |
| 1B | F0 | 0.959 | 0.201 | 2.71E-04 |
| 1B | F1 | 0.8 | 0.369 | 3.38E-06 |
| 1B | F2 | 0.689 | 0.349 | 1.29E-08 |
| 1B | F3 | 0.595 | 0.258 | 6.12E-06 |
| 1B | F4 | 0.523 | 0.262 | 2.58E-08 |
| 1B | F5 | 0.431 | 0.267 | 0.01256 |
| 1B | F6 | 0.237 | 0.264 | 0.1577 |
| Not shown in figure | F7 | 0.158 | 0.163 | 0.7674 |
| 2C | Drice[RNAi] | 0.295 | 0.291 | 0.6774 |
| 2C | Dcp-1[RNAi] | 0.329 | 0.261 | 0.4359 |
| 2D | Low protein v high (control) | 0.163 | 0.173 | 0.575213 |
| 5A | maternal | 0.754 | 0.244 | 0.000182 |
| 5A | paternal | 0.425 | 0.29 | 0.001031 |
| S1D | Brood 1 | 0.81 | 0.335 | 3.12E-08 |
| S1D | Brood 2 | 0.33 | 0.273 | 0.1903 |
| S1E | Exposed (1 gen) v Unexposed | 0.769 | 0.164 | 1.08E-05 |
| S1E | Exposed (2 gen) v Exposed (1 gen) | 0.908 | - | 1.82E-04 |
| S1E | Exposed (2 gen) v Unexposed | - | - | 1.82E-04 |
| S1F | Exposed (1 gen) v Unexposed | 0.737 | 0.172 | 1.82E-04 |
| S1F | Exposed F8 (2 gen) v Exposed (1 gen) | 0.784 | - | 0.08873 |
| S1F | Exposed F8 (2 gen) v Unexposed | - | - | 1.82E-04 |
| S2A | paternal (F1) | 0.569 | 194 | 1.08E-05 |
| S2B | Two-week old F1 | 0.833 | 0.127 | 0.000022 |

**Table S4.**
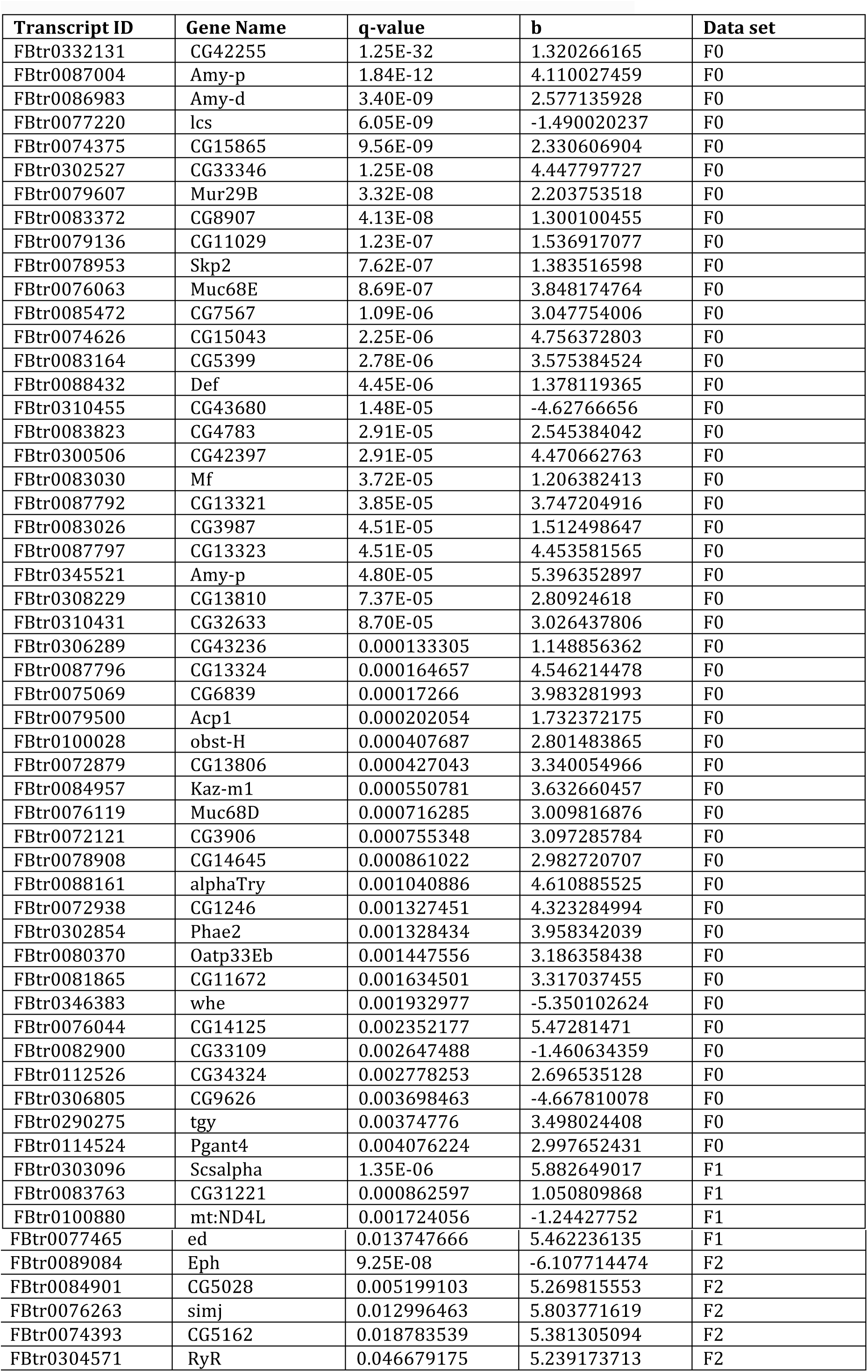
RNA sequencing results from female fly heads.

**Table S5.** Drosophila stock list and source information.

| name | genotype | source | stock number |
| --- | --- | --- | --- |
| CS | + | Bosco Lab |  |
| OreR | + | Bosco Lab |  |
| Orb2[deltaQ] | Orb2[deltaQ] | Bosco Lab |  |
| NPF-Gal4 | y[1] w[*]; P{w[+mC]=NPF-GAL4.1}2 | Bloomington Stock C | 25681 |
| UAS-NPF | UAS-NPF | Shen Lab |  |
| Elav-Gal4 | Elav-Gal4 | Bosco Lab |  |
| UAS-NPF[RNAi] | y[1] v[1]; P{y[+t7.7] v[+t1.8]=TRiP.JF02555}attP2 | Bloomington Stock C | 27237 |
| UAS-NPFR[RNAi] | y[1] v[1]; P{y[+t7.7] v[+t1.8]=TRiP.JF01959}attP2 | Bloomington Stock C | 25939 |
| UAS-Dcp1[RNAi] | y[1] v[1]; P{y[+t7.7] v[+t1.8]=TRiP.HM05120}attP2 | Bloomington Stock C | 28909 |
| UAS-Drice[RNAi] | y[1] sc[*] v[1]; P{y[+t7.7] v[+t1.8]=TRiP.HMS00398}attP2 | Bloomington Stock C | 32403 |
| Matα-Gal4 | w[*]; P{w[+mC]=matalpha4-GAL-VP16}V37 | Bloomington Stock C | 7063 |
| ninaB | w[*]/Dp(1;Y)y[+]; ninaB[1], P{w[+mC]=UAS-ninaB.G}3 | Bloomington Stock C | 24776 |
| compound ch-II | C(2)EN, b[1] pr[1] | Bloomington Stock C | 1112 |
| compound ch-II | C(2)EN, bw[1] sp[1] | Bloomington Stock C | 1020 |
| compound ch-III | C(3)EN, Diap1[1] st[1] | Bloomington Stock C | 1114 |
| compound ch-III | C(3)EN, st[1] cu[1] e[s] | Bloomington Stock C | 1117 |
| Df(3)10642 | w[1118]; Df(3R)ED10642, P{3'.RS5+3.3'}ED10642/TM6C, Sb[1] | Bloomington Stock C | 9482 |
| Df(3)BSC472 | w[1118]; Df(3R)BSC472/TM6C, Sb[1] cu[1] | Bloomington Stock C | 24976 |
| Df(3)BSC510 | w[1118]; Df(3R)BSC510/TM6C, Sb[1] cu[1] | Bloomington Stock C | 25014 |
| yw | y[1]w[1] | Bloomington Stock C | 1495 |

